# Use of null models to compare the assembly of northeast Atlantic bacterial community in the presence of crude oil with either chemical dispersant or biosurfactant

**DOI:** 10.1101/2020.12.23.424141

**Authors:** Christina Nikolova, Umer Zeeshan Ijaz, Tony Gutierrez

**Author notes:** **Corresponding author:** Dr Umer Zeeshan Ijaz, E.

## Abstract

The compositions of marine microbial communities in response to crude oil in the presence of biosurfactant or synthetic dispersants have been extensively studied in the last decade. Assembly processes, however, in such communities are poorly understood. In this study, we used seven different but complementing null model approaches, such as elements of metacommunity structure, Raup-Crick beta-diversity, normalised stochasticity ratio, Tucker’s null model, quantitative process estimates, lottery assembly, and phylogenetic dispersion models, to quantify the relative importance of ecological process that drive the community assembly. We found that the presence of chemical dispersant in the oil-amended microcosms induced significant temporal changes in the assembly processes that were different from the oil-only or biogenic dispersant-amended microcosms. The assembly processes in all microcosms were neither purely deterministic nor stochastic, but increasingly deterministic in dispersant-amended microcosms. Furthermore, the relative importance of determinisms varied over time and was strongest during the middle phase of incubation. Tucker’s null model revealed that phylogenetically distinct taxa might have shaped the bacterial community assembly in the different microcosms towards more niche or neutral processes. Moreover, there was faster recruitment of phylogenetically distant species in the dispersant-amended community. Drift, homogenising selection and dispersal limitation were the dominant assembly processes in all microcosms, but variable selection was only important in dispersant-amended microcosms. In conclusion, our study highlights that the assembly processes in marine bacterial communities are not static but rather dynamic, and the chemical dispersant can cause significantly different patterns of community assembly compared to non-amended or biosurfactant-amended microcosms.

**Importance:** The null model strategy is designed to intentionally exclude an ecological or evolutionary process of interest and create a beta diversity pattern that would be expected in the absence of this particular process – i.e. the community structure is random in respect to the process being tested. Recent advancements of bioinformatics and statistical tools have made it possible to apply theoretical macroecological concepts to microbial metagenomics in order to better understand and quantify the mechanisms and patterns controlling the complexity of microbial ecology. The conclusions from the null models can help predict the changes in microbial biodiversity and ecosystem services in oil polluted environments and therefore assist in making effective decisions with regards to what would be the best oil spill response option for similar environmental conditions.

## Introduction

Traditionally, community assembly is thought to be influenced by either stochastic (ecological births, deaths, extinctions, speciation, and colonisation) or deterministic (species traits, interspecies interactions, and environmental conditions) processes. Stochastic neutral theory assumes that species respond to chance colonisation, extinction, and ecological drift not because of their traits or that of their competitors, but rather because of random changes and thus have no interspecific trade-offs (i.e. species are neutral) (1). Niche theory, on the other hand, assumes that site-to-site variations in species composition is determined entirely by their specific traits, local habitat conditions, and interspecific relations, which in turn creates different niches that benefit different groups of species (determinism). The work of Chase and co-workers, however, stipulates that the assembly of local communities is simultaneously driven by both deterministic and stochastic processes (2, 3). Their framework is built around the concept that beta diversity patterns can resolve the relative influence of deterministic from stochastic processes in assembling the community structure along environmental gradients (e.g. space and time) (3). For this, related but different null model approaches can be applied to determine the relative importance that deterministic and stochastic processes have in generating patterns or variations in biodiversity.

Some null models use incidence-based (presence/absence of microbial species) metrics to calculate the null deviation between the observed and expected beta diversity (4–6) with the view that one can reorder the patterns (species and samples) to reveal distinct pattens that have ecological relevance. However, abundance-based data contain more information about species associations and distribution than the incidence-based data, and, therefore, deemed to be more reliable for understanding what are the underlying community assembly processes (7). For this reason the latest developed null model frameworks are built on abundance-based metrics (8) with phylogenetic information (9–12).

The use of technical jargon to describe ecological processes varies in scientific literature. Although some studies have been interpreting beta-null deviation values of zero as both, “stochastic” and “neutral” assembly, and deviation values different from zero as both “niche” and “deterministic” (5, 6, 13, 14), others pointed out that the terms do not represent the same thing (9, 15). While “neutrality” implies to a community in which species are equal and so share analogous demographic rates (birth, death, migration, etc.) (1), “stochasticity”, on the other hand, implies that the variation at which these demographic rates occur is random without implying anything about the mean values of these demographic rates (15). “Niche” entails the differentiation in mean demographic rates between species (16) and “determinisms” is the absence of random variations in species’ demographic rates (9). In fact, Tucker et al. (9) built a new computational framework that allows for more appropriate interpretation of the beta-null deviation measure – i.e., differentiation between neutral and niche community. According to their model, beta-null deviation values close to zero indicate neutral assembly, and values that are different from zero as niche-based assembly. Furthermore, Tucker et al.’s framework has been applied in a study that investigated the response of soil microbial communities to an anthropogenic disturbance (10) and in another that explored the role of ecological selection, drift and dispersal in assembling the bacterial communities associated with domesticated and wild wheat species (17).

Ecological process can be used to characterise the distribution of species in the microbial community assembly and identify groups or guilds of phylogenetically related species whose distribution may be governed by strong priority effects (i.e. competitive lottery model; see (18). A lottery-based assembly model was developed by Verster and Borenstein (19) to test to what extent the competitive lottery model applied to the human gut microbiome. The authors postulated that lottery-based assembly model could explain the variation in observed diversity of human gut microbial species between different hosts. This model assumes that there is a “winner” species (hence the name “lottery”), determined randomly, that solely occupies a given niche due to a competitive advantage over other species (18). Although the lottery winner provides information on whether the distribution of phylogenetically related species is governed by priority effects, the order in which new species are recruited in the ecosystem remains to be described. To tackle this, Darcy et al. (12) developed a null model-based framework to better understand how phylogenetic relationships between microbes influence the order in which microbes are recruited over time, and whether this recruitment occurs slow or fast. The authors applied their model to longitudinal human microbiome data which showed that human microbiome generally follows what they called the “nepotism” hypothesis, i.e., close relatives are more likely to be recruited in a community than distant relatives (also known as phylogenetic underdispersion) (12). This observation is in contrast to the expectation that closely related species would experience strong competition between each other so that they cannot coexist, but hint to growing support that phylogenetic underdispersion may be more common trend in microbial communities (20).

In this study, we used seven different but complementary null models to better understand and quantify the relative importance of the different assembly processes on a temporal scale in a Northeast Atlantic marine microbial community amended with crude oil dispersed with either chemical dispersant Finasol or rhamnolipid biosurfactant (21). To the best of the author’s knowledge, this study presents the first comprehensive study to apply all seven null models to oil-amended marine microbial communities to better understand the ecological processes behind the community assembly, and if and how the presence of an oil-dispersing agent (i.e., a biosurfactant and/or a synthetic chemical dispersant) had an effect on the assembly.

## Results

### Elements of metacommunity structure (EMS)

All treatments showed significantly negative coherence z-values indicating that microbial communities in the treatments were shaped by distribution patterns of competitive exclusion – i.e., determinism (Fig. 1). The crude oil-amended treatments with the addition of either Finasol (CEWAF) or rhamnolipid (BEWAF) did not show wide variation between each other’s coherence z-values but were lower than in the WAF treatment. However, the CEWAF treatment displayed the highest boundary clumping as denoted by the Morisita’s index. The *in-situ* FSC community had the lowest coherence z-value with Morisita’s index of 1, indicating that the spatial distribution of species in this metacommunity is over-dispersed. In the opposite end was the untreated seawater (SW) control treatment which had the highest coherence z-value but also the highest negative turnover z-value (−8.56). The seawater treatment amended with Finasol only (SWD) also had negative turnover z-value (−5.45) indicating that the species distribution is caused by species loss.

**Fig. 1.**
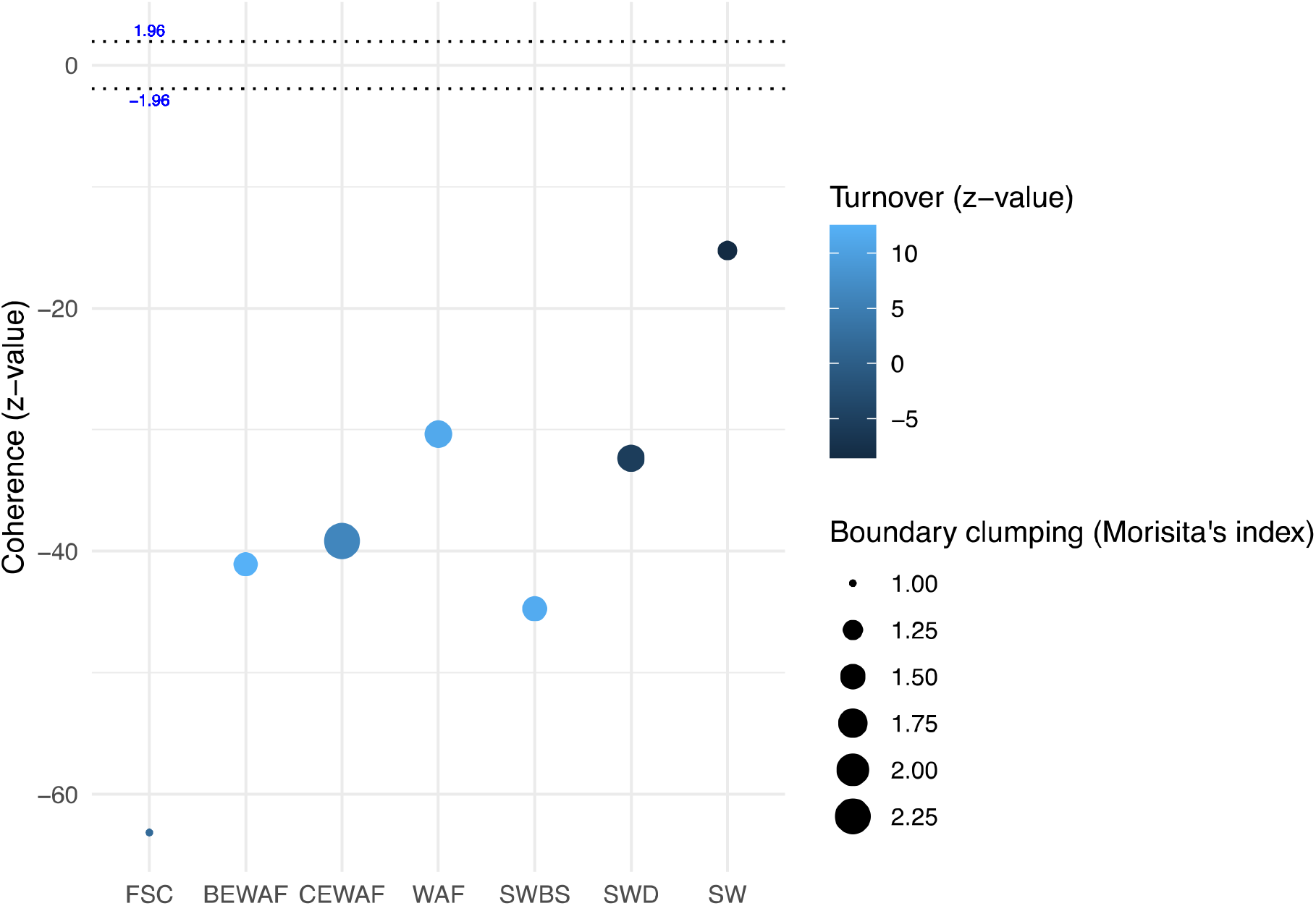
Variation of metacommunity types of the bacterial 16S rRNA sequences shown for each treatment. The blue scale bar represents species turnover (z-value; number of observed replacements compared to a null distribution) where positive values indicate species replacements in response to environmental variation and negative values nested species distributions caused by special losses. The size of the circles denotes the Morisita’s index (boundary clumping) which shows the degree of spatial distribution of species in a metacommunity where lower numbers indicate over-dispersed boundaries and higher numbers clumped boundaries (analogous to clustering of microbial species). Metacommunities are randomly structured when −1.96 < coherence z-value > 1.96. Positive significant values (coherence z-value >1.96) indicate that species contribution occurs in response to environmental variation. Significantly negative coherence (coherence z-value < −1.96) indicates competitive exclusion distribution.

### Incidence-based beta diversity (β_RC_)

The β_RC_ dissimilarity indices were calculated to test whether communities in each treatment were assembled due to stochastic or deterministic processes. In general, all treatments, except the *in-situ* FSC community, varied within a narrow range, not deviating strongly from – 0.5, which indicates that the processes responsible for the community assembly in each treatment could be neither strongly stochastic nor deterministic. However, the microbial communities seemed more similar to each other than expected by chance (Fig. 2). The *in-situ* FSC community did not deviate strongly from the null expectations (β_RC_ = 0.166) and, therefore, the microbial community was more likely to be assembled by stochastic processes.

**Fig. 2.**
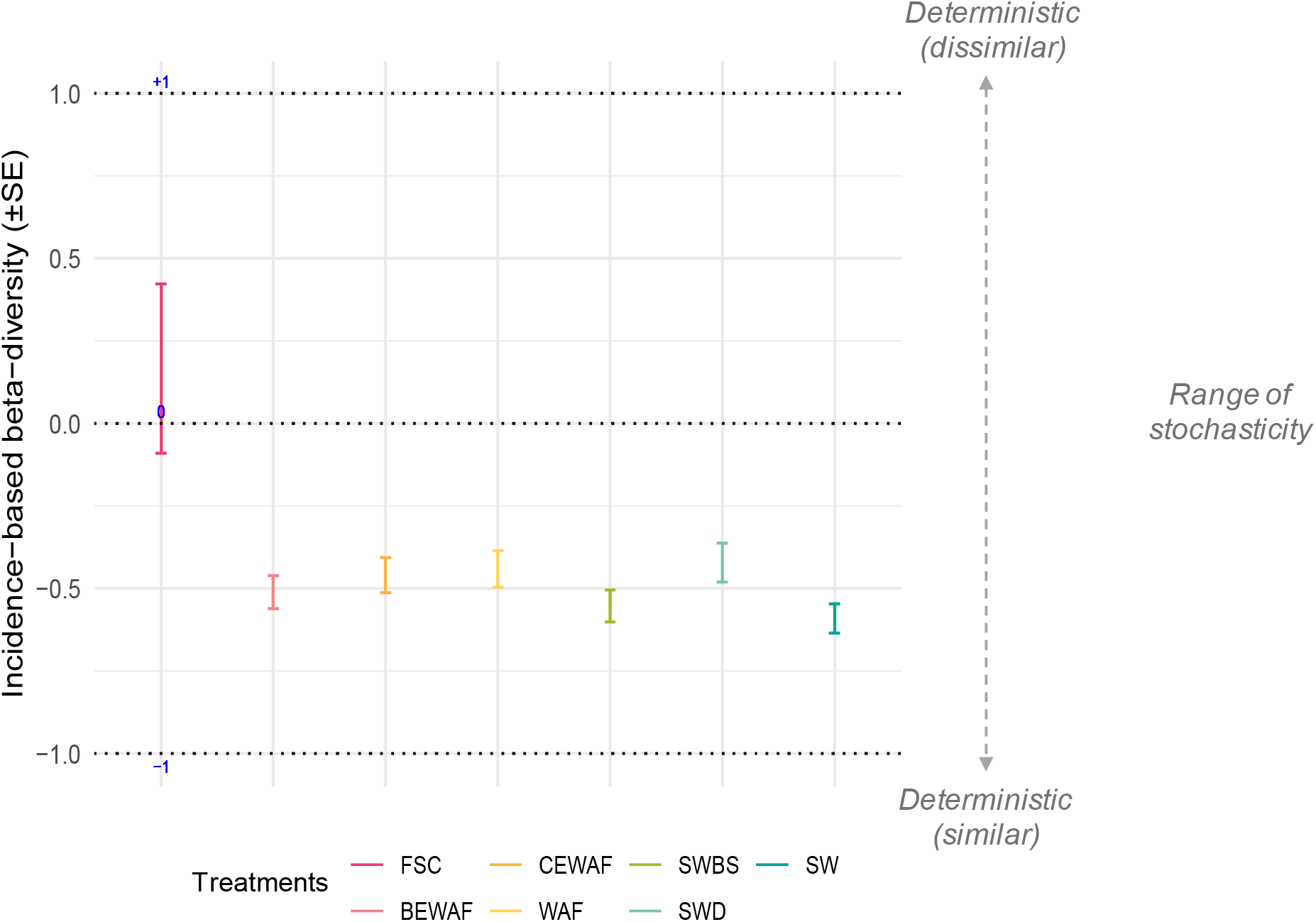
Variation of incidence-based (Raup-Crick) beta diversity (β_RC_) for *in-situ* seawater (FSC), crude oil amended seawater (WAF), crude oil + dispersant amended seawater (CEWAF), crude oil + rhamnolipid amended seawater (BEWAF), seawater amended with Finasol (SWD) or rhamnolipid (SWBS), and non-treated seawater (SW; control) treatments.

### Stochastic vs. deterministic assembly processes on a temporal scale

In general, the NST revealed that the microbial community assembly in the studied treatments were neither purely deterministic nor purely stochastic as indicated in Fig. 3A. The *in-situ* FSC community, the crude oil and seawater WAF treatment, and the seawater only control (SW) had the same NST value of 56%, while the rhamnolipid-amended oiled seawater treatment (BEWAF) had a NST of 51%, suggesting that the communities’ assembly in these treatments were driven predominantly by stochastic processes. Communities in the treatments with added chemical dispersant Finasol (CEWAF and SWD), in contrast, were dominated by deterministic processes as their NST was, respectively, 38% and 35%. PERMANOVA analysis revealed that the CEWAF treatment was significantly different from the SW control treatment *(p* = 0.019) and the SWD treatments was significantly different from the BEWAF *(p* = 0.038), WAF *(p* = 0.018), and the SW *(p* = 0.002) treatments. The rest of the treatments were not found to be significantly different from each other (**Error! Reference source not found.**).

**Fig. 3.**
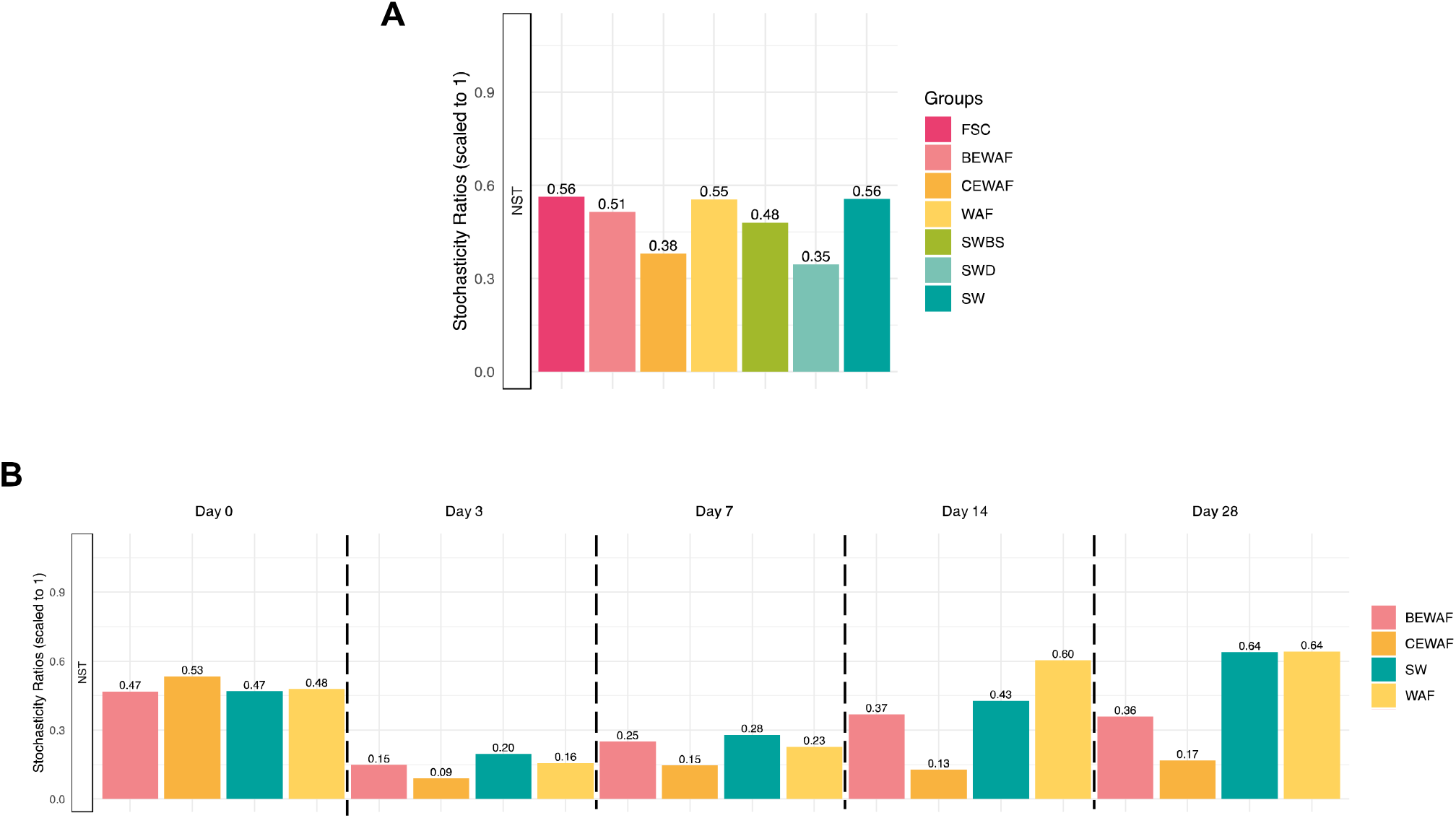
(A) Normalised stochasticity ratio (NST) for *in-situ* seawater (FSC), crude oil amended seawater (WAF), crude oil + dispersant amended seawater (CEWAF), crude oil + rhamnolipid amended seawater (BEWAF), seawater amended with Finasol (SWD) or rhamnolipid (SWBS), and non-treated seawater (SW; control) treatments. NST was calculated based on abundance based Ružička metric using null model algorithm PF which stipulates that the probabilities of taxa occurrence are proportional to the observed occurrence frequencies, and taxon richness in each sample is fixed. (B) Temporal changes of the estimated NST. NST was calculated based on abundance-based Ružička metrics using null model algorithm PF.

Next, the NST was determined for each treatment in respect to temporal dynamics The NST method becomes less precise when there are less than 6 replicates (6), therefore, the results presented in Fig. 3B should not be taken as absolute true but rather as a guidance. Because of not enough replicates per treatment (n ≤ 3) at each time point, PERMANOVA was not performed. Overall, stochasticity varied substantially over time in all treatments. On day 0, all presented treatments had similar NST values close to the 50% boundary point (45 – 53%). However, by day 3 the NST dramatically dropped in all treatments to 16% (WAF), 15% (BEWAF), 9% (CEWAF), and 20% (SW) suggesting that the microbial communities were overwhelmingly driven by deterministic processes and even more so in the CEWAF treatment. From this point forward, the estimated NST started to increase in all treatment but was lowest in the CEWAF treatments until the end of the incubation period. Stochasticity was high at day 28 for the WAF and SW treatment (over 60%) but remained low in the BEWAF and CEWAF treatments. Collectively, these results indicate that stochastic processes could play more important roles in controlling community succession in its early and late phases, while deterministic processes could be more important during the middle phase.

### Tucker’s beta null model

To quantify the relative contribution of niche and neutral processes in the assembly of the bacterial communities in seawater microcosms amended with crude oil (WAF) and dispersant (CEWAF) or rhamnolipid (BEWAF), Bray-Curtis and weighted UniFrac beta-diversity deviations from null expectation were computed. The Bray-Curtis (abundance only) and weighted UniFrac (abundance and phylogenetic relatedness) beta-null deviation results showed a dynamic pattern for the three treatments over time (Fig. 4). The WAF treatment deviated from the null expectation towards a significant increase in the contribution of niche processes in the community assembly by the end of the incubation time period. In contrast, the relative contribution of niche processes in the BEWAF and CEWAF treatments varied more over time. The assembly of communities in both treatments was more prone to neutral processes during the first 3 days of incubations. Afterwards, the beta null deviation in the CEWAF treatment transitioned to more niche processes, before going back to neutral processes on day 14, and finally to an increase in niche processes’ relative contribution. The analysis of the weighted UniFrac deviation from null model indicated that the relative contribution of neutral processes was more dominant in the BEWAF treatments assembly over the late phase of the studied period (day 14-28) in contrast to the Bray-Curtis null deviation (Fig. 4). BEWAF’s Bray-Curtis null deviation was not significantly different from that of the CEWAF on day 28, but it was statistically different when taking into account the phylogenetic relatedness (weighted UniFrac), indicating that phylogenetically distinct ASVs between the two treatments might play a role in shaping the bacterial community assembly towards more niche or more neutral processes.

**Fig. 4.**
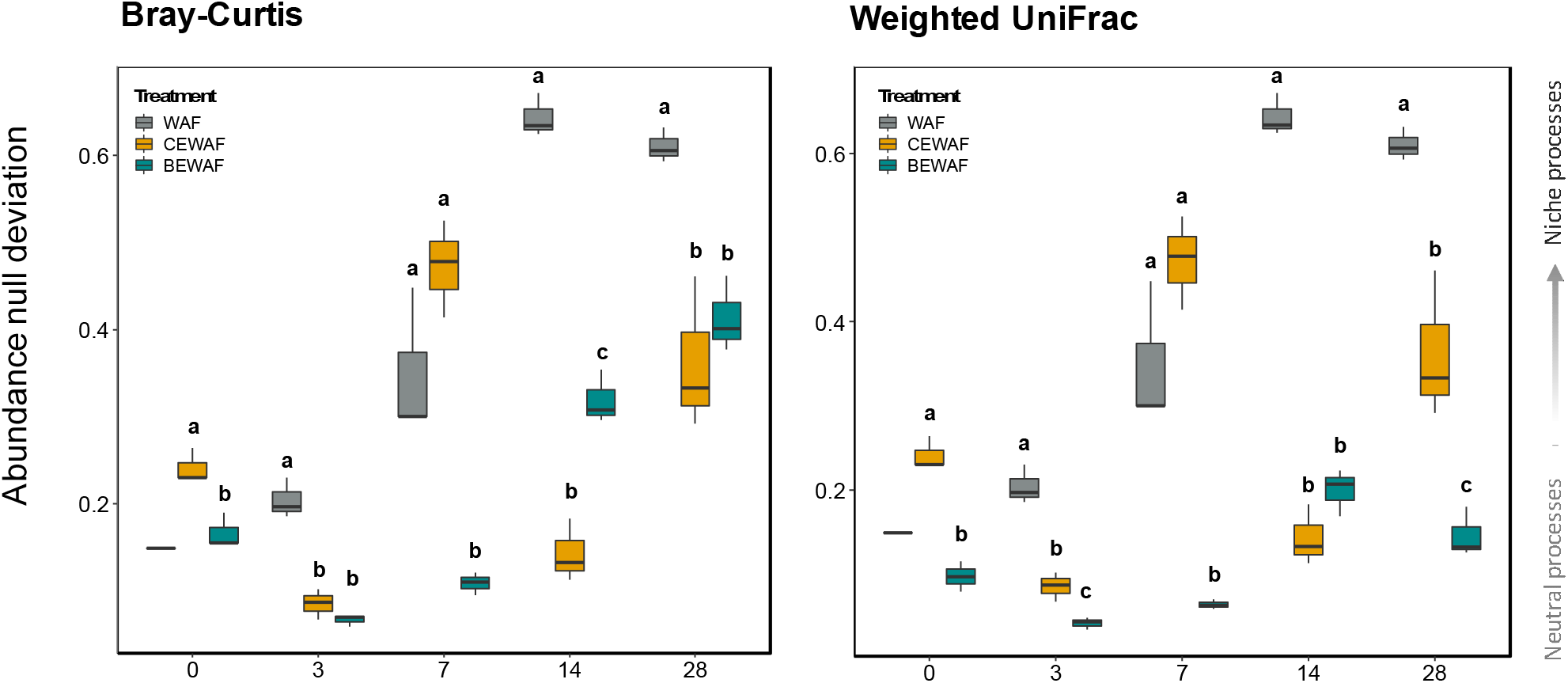
Boxplots of Bray-Curtis (left) and weighted UniFrac (right) abundance deviation from the null model of the microbial communities in treatments WAF, CEWAF and BEWAF over time (days 0 to 28). Significance between group means for each time point were tested using Two-way ANOVA analysis and *post-hoc* Tukey’s test. Groups that share different letters are significantly different from each other.

### Quantitative process estimates (QPE)

The aim of the QPE was the estimate the relative importance of assembly processes. Considering the overall dynamics of these processes in the different treatments, there are some clear differences. For the *in-situ* FSC community, random processes or drift was the dominant assembly process (67%) followed by homogenising selection (33%) (Fig. 5). On the other hand, in all treatments, the dominant assembly processes were ecological drift, homogenising selection and dispersal limitation. The variation in the relative proportion of ecological drift among the treatments was greater than the variation in the rest of the assembly processes. Ecological drift was the least important in the CEWAF (29.5%) and the SWD (35.9%) treatments, while in the WAF and seawater control (SW) treatments the drift accounted for more than 50% of the assembly processes (Fig. 5). Homogenising selection had similar relative importance in all of the treatments (except the *in-situ* FSC) ranging from 19.7% in the WAF to 27.6% in the CEWAF treatment. Dispersal limitation was also similarly important in the assembly of communities in all treatments (20.9-27.6%) except for the *in-situ* FSC community and the SWD treatment where the dispersal limitation was not important at all (0%) or in very small proportion compared to other treatments (3.8%), respectively. Among all treatments, variable selection was relatively important in only two treatments – CEWAF (17%) and SWD (30.7%), while for the other treatments it was not found to be relatively important. The ecological process that had the lowest relative influence in the community assembly among all treatments was the homogenising dispersal. In fact, it had less than 1% of effect in the treatments BEWAF, CEWAF, SWBS and SW, just above 1% in the CEWAF and was highest in the SWD treatment (7.7%).

**Fig. 5.**
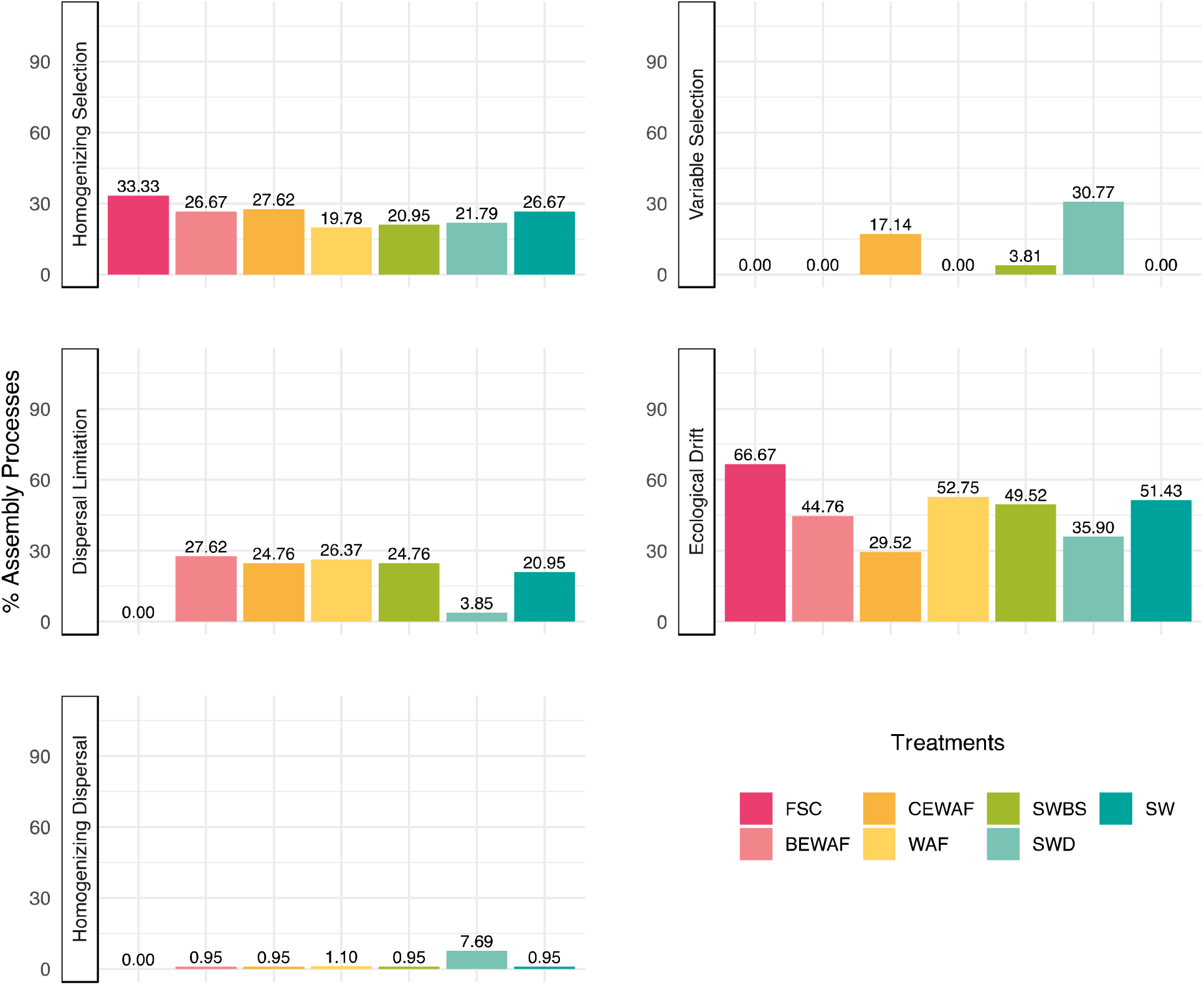
Overall dynamics of the relative importance of different community assembly processes expressed as the proportion of community pairs assembled either by species-sorting (variable or homogeneous selection), dispersal limitation or historical contingency, homogenising dispersal or ecological drift.

### Competitive lottery-controlled genera

The distribution of species across treatments and identity of lottery “winners” were characterised with the help of the lottery-based assembly model. The ASV distribution was expected to display two fundamental features. First, a single group member captures > 90% of the group’s abundance (the “lottery winner”) in each treatment and, second, different treatments should have different lottery winners. The winner prevalence (the fraction of samples in which one ASV was assigned > 90% of the genus abundance) and the winner diversity (the normalised diversity of lottery winners) for each genus in the WAF, CEWAF and BEWAF treatments were plotted in Fig. 6. There was a number of genera with high winner prevalence. For example, *Pseudophaeobacter* was the lottery winner in the WAF treatment as it was present in 85% of samples and showed highest winner diversity (41%). Similarly, in one ASV of *Cycloclasticus* was in 71% of the WAF samples but it had a higher winner diversity than *Pseudophaeobacter.* The lottery winner in the BEWAF treatment was *Pseudohongiella* which was present in 80% of the samples. In the CEWAF treatment, the genus with the highest winner prevalence was a different *Cycloclasticus* ASV (from the one in the WAF treatment) but it had a winner diversity of 1, meaning that this particular *Cycloclasticus* ASV was not consistent with the competitive lottery schema. Groups that have low winner prevalence and have comparatively higher winner diversity likely reflected that the group abundances were more evenly distributed among the group ASVs (e.g., *Peredibacter, Hyphomonas).*

**Fig. 6.**
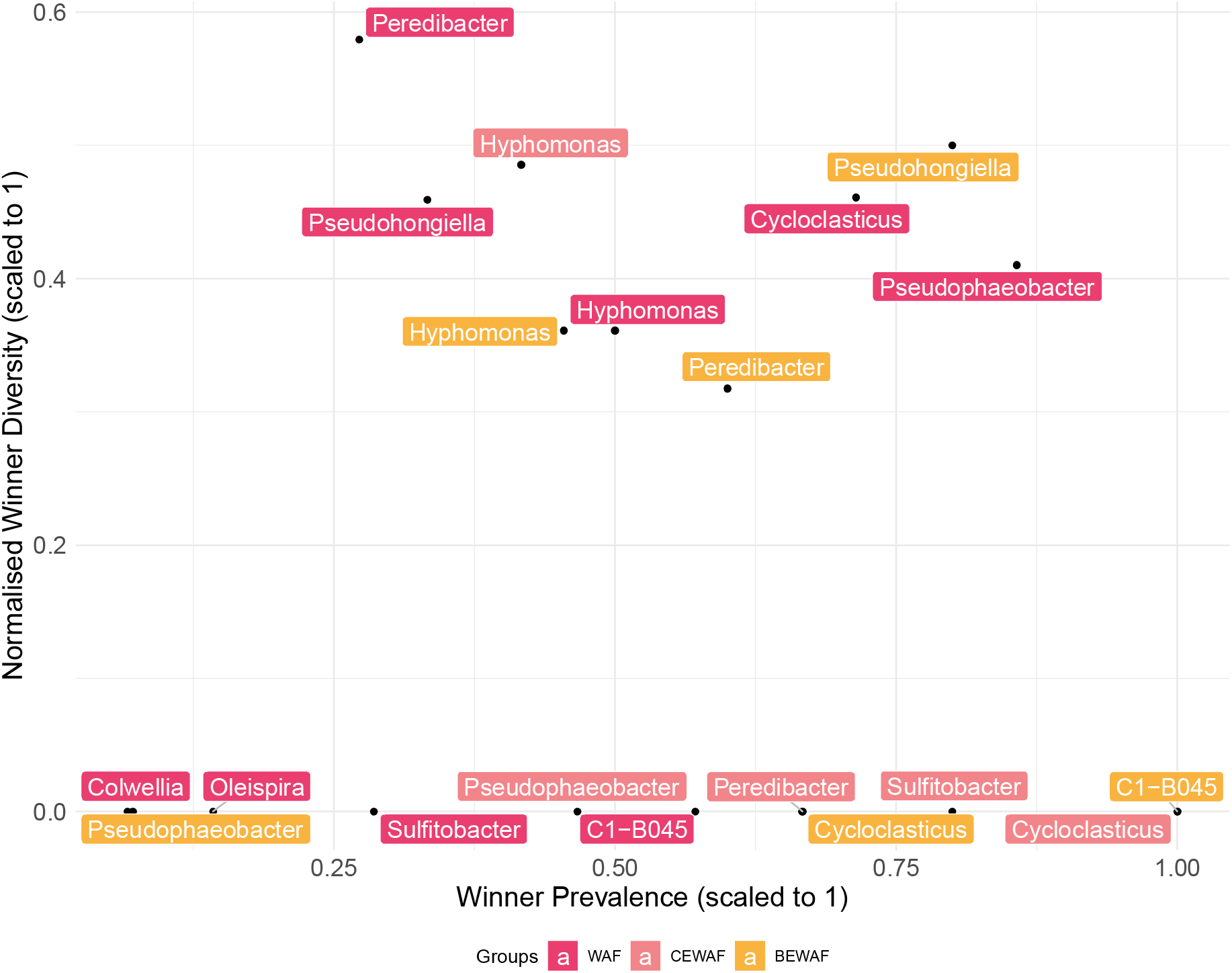
A scatter plot showing the winner prevalence and winner diversity for different genera in three crude oil-amended seawater treatments: WAF (oil + seawater), CEWAF (oil + seawater + Finasol), and BEWAF (oil + seawater + rhamnolipid).

### Phylogenetic dispersion

The phylogenetic dispersion null model was used to estimate the extent of recruitment of new species into the microbial community of six natural seawater microcosms (Fig. 7). Varying 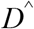 changed the rate at which the phylodiversity was added to the resampled microbial communities over time and overall, the results showed that the *D* parameter successfully corresponded to over- and underdispersion relative to the neutral model (Fig. S1). The CEWAF treatment had the highest *D* value (*D* > 0) compared to the other two oil-amended oils (WAF and BEWAF), indicating that the presence of dispersant changed the phylogenetic colonisation patterns to a preferential and faster recruitment of phylogenetically distant species in the community (i.e., overdispersion) (Fig. 7). In contrast, the WAF and BEWAF treatments had *D* value of less than 0 which suggests that there was underdispersion or phylogenetically similar new species were detected in the local community. In other words, the addition of dispersant caused the local community to become phylogenetically divergent as time progressed, whereas the addition of rhamnolipid caused the opposite trend – a phylogenetically constrained community. The oil by itself (WAF) also caused constrained community but less so compared to the rhamnolipid-amended oil treatment (BEWAF). All treatments had significantly different *D* values *(p* < 0.001) (Fig. 7).

**Fig. 7.**
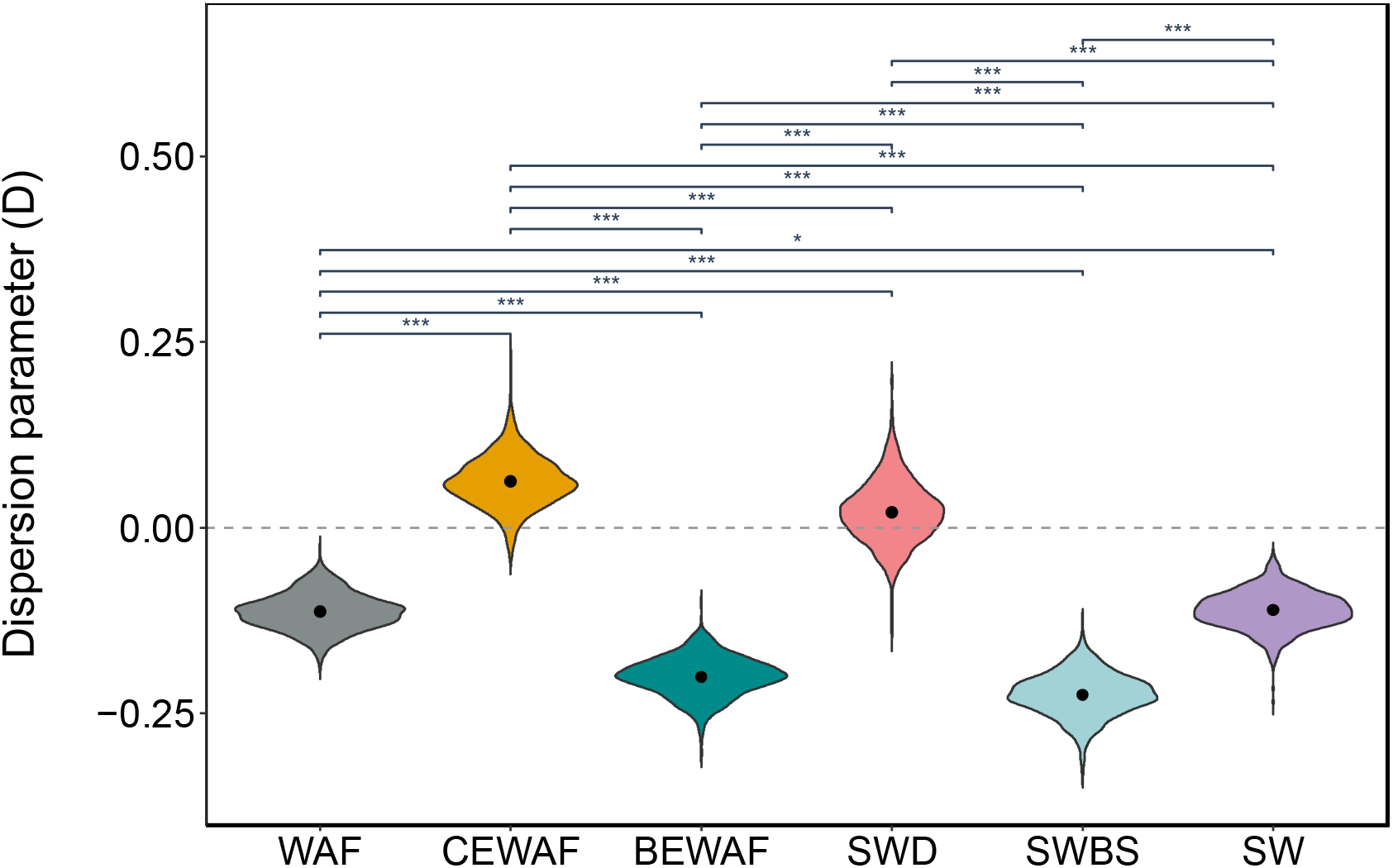
Violin plot of the distribution of dispersion parameter (*D*) estimates given by logistic error model bootstrap for six treatments (fill colour). Dots within each violin are means. Treatments are seawater (SW) amended with crude oil (WAF), crude oil and dispersant Finasol (CEWAF), crude oil and rhamnolipid biosurfactant (BEWAF), Finasol only (SWD), or rhamnolipid only (SWBS) over time. Significantly different (t.test) treatments are share a bracket and the level of significance is shown with * *(p* = 0.05), ** *(p* = 0.01), or *** *(p* = 0.001).

## Discussion

Seven different and recently developed null model frameworks were used to quantify the relative importance of ecological processes in the community assembly (3, 4, 6, 9–12, 22). Elements of metacommunity structure (EMS), Incidence-based (Raup-Crick) beta-diversity, and normalised stochasticity ratio) were developed to detect patterns in binary incidence (i.e., presence/absence) matrices taking into account only taxonomic beta-diversity estimates. The next model used in this study was the beta-null framework developed by Tucker et al. (9) to differentiate between neutral and niche communities and was based on quantitative abundance-based (Bray-Curtis) dissimilarity matrix. It was later complemented by Lee et al. (10) to integrate phylogenetic relatedness information with the taxonomic abundance (weighted UniFrac). The QPE framework was also based on abundance and phylogenetic-based matrices to quantify the relative importance of ecological selection, drift, and dispersal. Although the lottery-based assembly model used in this study is abundance-based, it focuses more on the characterisation of the species distribution across treatments in relation to priority effects (11). The phylogenetic dispersion model provided a view into temporal dynamics in the recruitment of new species in the local community in relation to their phylogenetic similarity or dissimilarity to already existing members of the community which colonised it at a previous time point (12).

### Stochastic vs. deterministic assembly

It has been generally accepted that both deterministic and stochastic processes occur simultaneously in the assembly of local communities (5). According to incident-based betadiversity (β_RC_) patterns, bacterial communities across all communities studied here were neither purely stochastic nor purely deterministic. However, there was a tendency toward more deterministically assembled communities (β_RC_ ~ −0.5). EMS provided further evidence for deterministic assembly, and the NST ratio supported this finding. In particular, the NST showed that the oil-only control treatment (WAF) and untreated seawater control (SW) had more stochastically assembled communities, while the communities in the treatments with added dispersant Finasol (CEWAF and SWD) were more deterministic. It is likely that the presence of dispersant triggered a microbial response related to deterministic succession. Furthermore, NST also revealed that the relative importance of stochasticity over determinism varied substantially over time across treatments. Stochastic processes were more prevalent in the early and late phases of incubation, while deterministic processes are more important in the middle phase in all treatments. The result in the middle phase seems fit to intuition that adding carbon source (e.g., crude oil and/or dispersant/biosurfactant) or altering the environmental conditions should drive selection and hence leads to a more deterministic outcome. This is a demonstration that drivers controlling biodiversity and community succession are dynamic rather than static in fluidic ecosystems (23, 24).

### Neutral vs. niche assembly

The Tucker’s beta-null deviation model successfully differentiated patterns of niche and neutral processes in the three crude oil amended treatments (WAF, CEWAF, and BEWAF) in the presence of either dispersant or rhamnolipid over time, suggesting that the presence of dispersant, either synthetic (Finasol) or biogenic (rhamnolipid) did have an important role or even were selective factors in the community assembly processes. Neutral processes had a more prominent role in the assembly in the rhamnolipid-amended oil treatments than in the oil-only and Finasol-amended oil treatments. This finding suggests that the addition of rhamnolipid had not applied strong selection on the assembly of oiled seawater microcosms. The opposite was observed for the oil-only treatment (WAF) – clear niche community assembly pattern. A plausible explanation is that the oil selected for highly specialised species while the addition of rhamnolipid allowed more generalist taxa to thrive (see 21).

### Importance of selection, dispersal and drift

The microbial succession patterns in response to either dispersed or non-dispersed crude oil in the marine environments have been well documented (25–31). In all cases, there was a distinct succession of obligate and generalist hydrocarbon-degrading taxa which rapidly increased in abundance to dominate the community structure composition. Furthermore, the microbial respond to chemically dispersed oil has been shown to be different to oil-only treatments (32, 33). But which ecological process and to what extent do they govern such a response has not been explored before, probably due to the many uncertainties around microbial ecology and/or the significant computational power required to process large 16S rRNA sequencing data.

The QPE framework showed that all treatments were dominated by drift, dispersal limitation, and homogenising selection. Homogenising dispersal was not found to be an important factor in the community assembly as expected for a marine ecosystem where there is a rapid dispersion and population movement (i.e., high connectivity) (23, 34). Drift was the most dominant ecological process detected in the QPE analysis in the majority of treatment. The importance of drift was notably lower in the both Finasol-amended treatments (CEWAF and SWD) compared to the rest of the treatments. The dispersal limitation was relatively higher than it would be expected for open ocean ecosystem likely because the microbial microcosms in this study (see 21) were enclosed in bottles where there was no water exchange, although constant rotation was applied to ensure adequate mixing and simulate the motion of seawater in the upper water column. The relative importance of dispersal limitation in the *in-situ* water sample (FSC), in comparison, was 0%, thus supporting that dispersal limitation in open ocean systems is not a decisive factor in community assembly. There is growing evidence, however, to support that the dispersal limitation can actually be a more important dominating factor in marine microbial communities than previously thought, especially when acting together with drift (35, 36). Other microbial studies using quantitative process estimates have demonstrated the substantial proportion of the dispersal limitation or historical contingency (such as priority effects) in community assembly (37–39). It is reasonable to suggest that in a real-life oil spill, dispersant or biosurfactant application would not have a strong deterministic effect due to the higher dispersal rates observed in oceanic systems. The patterns of dispersal limitation and homogenising dispersal in the dispersant-only control treatment (SWD) particularly stand out from the rest of the treatments. It seems that the dispersant itself had high relative influence over the ecological processes. Indeed, the importance of variable selection was substantially higher only in the treatment containing Finasol dispersant (CEWAF and SWD). One explanation for this could be attributed to a smaller number of taxa that took over the community structure in CEWAF (21) that otherwise be selected against in communities without dispersant because of spatial variation in the selective environment. In fact, the variable selection causes an increase in the spatial environmental heterogeneity as time (i.e., succession) progresses which leads to compositional differences across local communities (24). Moreover, regression analysis confirmed that the dispersant caused a reduction in alpha diversity (21). Generally, selection has more detectable influence over microbial communities (6–8) and the same was observed in this study. Relatively strong homogenous selection was expected as selection is likely to be consistent across spatially homogenous systems (23), such as the natural surface seawater ecosystem studied here. Stochasticity is influenced by an interaction between dispersal and selection, with stronger selection causing an increase in drift or priority effects (40). The importance of drift is considered higher when selection is weak, and the local community is small. Pure drift is practically impossible to measure, and even more so for microbial communities, because no species in nature are exactly the same in a demographic sense. Distinct populations that share similar or the same ecological function (i.e., functional redundancy) are quite common in microbial communities. Functional redundancy tends to increase neutrality and sensitise functional redundant populations to ecological drift, which is unarguably stochastic. In conclusion, it was the direct or indirect interrelation of selection, dispersal, and drift with each other that assembled the microbial community structures in this study, as observed elsewhere (40).

### Lottery winners

The findings of the lottery-based assembly model presented in this study confirmed that the lottery winner ASVs varied between treatments in accordance with the competitive lottery schema. Furthermore, the lottery winners in each treatment did not occur at the same frequency as assumed by the lottery schema (11). A winner diversity approaching 0 represents that same species is selected in all cohorts for a given clade. There were a number of genera with very high winner prevalence but very low winner diversity, including *Cycloclasticus* and *Sulfitobacter*, that did not involve complete competition-derived exclusion but rather strong coexistence with other species in the microcosms or even more complex assembly that combines exclusion and coexisting patterns (11). Taking into consideration the relative abundance analysis of the top 25 most abundant ASVs observed in Nikolova et al. (21), it became apparent that the lottery winners in each treatment were not necessarily the most abundant species identified. For example, *Cycloclasticus* was identified as a potential lottery winner in the WAF treatment. Furthermore, *Cycloclasticus*, which is a PAH degrader, was found to dominate the WAF’s community composition in the late stages of incubation, while early stages were previously dominated by *Colwellia* and *Oleispira*, which are aliphatic and low-molecular-weight PAH degraders. It is logical then to assume that microbial succession, driven by competition for resources, occurred in the WAF treatment – i.e., *Cycloclasticus* outcompeted *Colwellia* and *Oleispira*. The relative abundance analysis performed in Nikolova et al. (21) showed that *Cycloclasticus* had a relative abundance of <1% across all CEWAF samples and it is highly likely that just one ASV was entirely responsible for the observed low abundance. According to the lottery-based assembly model, a single *Cycloclasticus* ASV was present in 100% of all samples in the CEWAF treatments (Fig.) but did not fit the assumptions of the competitive lottery schema. This suggests that this particular *Cycloclasticus* ASV was either outcompeted by other species (potentially more adept to the presence of dispersant or able to engage in resource partitioning) observed in the CEWAF treatment (e.g., by *Rhodobacteraceae)* to the point where its abundance never exceeded 1%, or the presence of the dispersant Finasol itself might have selected against it. Interestingly, generalist species *Pseudophaeobacter* and *Pseudohongiella* were projected the lottery winners in the WAF and BEWAF treatments, respectively, but both had less than 1 % abundance in the respective treatments (21). In fact, *Pseudophaeobacter* represented up to 40-fold higher abundance in the CEWAF treatment, while *Pseudohongiella* was most abundant in the WAF treatment. It is possible that there was other more complex assembly schema, which the lottery-based assembly method could not explain, that *Pseudophaeobacter* and *Pseudohongiella* conformed to in comparison to *Cycloclasticus*. For example, although they were lottery winners, both species could have facilitated subsequent species that join the ecosystem (i.e., respond to crude oil) to flourish. It is also possible that as generalists, *Pseudophaeobacter* and *Pseudohongiella* have more opportunities for niche diversification.

### Phylogenetic dispersion

The phylogenetic dispersion model is the most recently developed null model and hence, it has not been tested in other environmental microbial studies. Nevertheless, the model provides a valuable insight into the largely unknown area microbial ecology of how and when microbial communities are colonised by phylogenetically similar or dissimilar relatives (12). Microcosms treatments with added dispersant Finasol (CEWAF and control SWD) were the only two treatments that had *D* values > 0 indicating that the microbial community in these treatments become more phylogenetically divergent as time progresses and there is little preference of which species are recruited first (i.e., random colonisation). The rest of the treatments (WAF, BEWAF and the controls SWBS and SW) had *D* values < 0 and followed a “nepotistic” pattern of new species recruitment. This “nepotistic” pattern was described as a recruitment pattern in which new species that are closely related to the already existing member of the community are more likely to be recruited than distantly related species (12, 20). Traditionally, community ecology assumes that competition among closely related species would be strongest but also allows similar species to coexist, especially when dispersion is high as is in ocean ecosystems (20). One explanation for the overdispersion observed in the CEWAF treatment is that the dispersant caused the formation of multiple environmental gradients in the community which provided an opportunity for different species to colonise. This was further supported by the findings of the QPE model which showed that variable selection was only observed in the CEWAF and SWD treatments. Conversely, variable selection in BEWAF and WAF treatments was not detected, and hence, explain the observed phylogenetic under dispersion which posits that recruitment of new species is slow, i.e., there is a single environmental gradient available to colonise. This is somewhat opposite to the Tucker’s beta null model which showed that the community in the BEWAF treatment was assembled more by neutral processes than the CEWAF treatment.

In summary, our findings highlight that CEWAF treatment was more deterministic (environmental pressures) than the rest of the treatments, with the deterministic assembly processes dominating the middle phase of incubation. Application of numerous null models have revealed marked differences in microbial consortia structure between the dispersant, CEWAF and the biosurfactant amended treatment, BEWAF, and have unveiled insightful structures indicative of the underlying ecological processes. In view of our findings, it is also recommended to apply multitude of null-models and then find the consensus agreement or consistent patterns amongst them. This is particularly important as the methods capture different aspects of microbial consortia, whether focusing on abundances, or on phylogeny, and may also have analytical biases.

## Materials and Methods

### Sample collection, experimental set up, barcoded 16S rRNA amplification, and null modelling

Surface seawater sample was collected from the Faroe-Shetland Channel in the northeast Atlantic Ocean and along with the subsequent microbial microcosms set up, DNA extraction, and 16S rRNA amplicon Illumina sequencing used to perform the null models in this study have been described elsewhere in detail (21). The seven null models used in this study, i.e., elements of metacommunity structure (EMS), incidence-based (Raup-Crick) beta diversity, normalised stochasticity ratio (NST), Tucker’s beta-null model, quantitative process estimates (QPE), lottery-based assembly model and phylogenetic dispersion model, have been described in detail in Supplementary Information. All null models and visualisations were conducted in R version 3.5.3 (41).

## Acknowledgements

None

---

**Supplementary Table S1**. Output of PERMANOVA test for differences in NST among treatments. P.anova is the p-value of parametric ANOVA test; P.panova is the p-value of permutational ANOVA (PERMANOVA) test, and P.perm is the p-value of permutational test of the difference. Significantly different PERMANOVA values between treatments are shown in bold.

**Supplementary Figure S1**. Plots **a-c** and **g-i** show the empirical (dashed) and surrogate polydispersity accumulation curves coloured according to 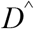 value. Blue colour corresponds to new species that have a previously detected close relative contribute little polydispersity and cause slow phylodiversity accumulation. Green colour corresponds to new species, that do not have a close relative, contribute more phylodiversity and cause faster accumulation. Teal colour corresponds to the neutral model which is above the empirical model (dashed line), signifying underdispersion in the order of first-time species detections. The time of sampling points (days) are shown on the x-axis. Plots **d-f** and **j-l** show how empirical and surrogate data compare with generation of *D* estimates. The y-axis represents the difference between empirical and surrogate data at time *m* and 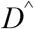 value used to generate the surrogate datasets is shown on the x-axis. Colour-coded points correspond to surrogate datasets, grey-coloured points are extreme values of 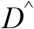 (which help to fit the logistic error model – black line), and the red arrow show the process of error minimisation, producing a *D* estimate.

## References

1. Hubbell SP. 2001. The unifed neutral theory of biodiversity and biogeograophy (MPB-32). Princeton University Press.

2. Chase JM. 2010. Stochastic community assembly causes higher biodiversity in more productive environments. Science 328:1388–1391.

3. Chase JM, Myers JA. 2011. Disentangling the importance of ecological niches from stochastic processes across scales. Philosophical Transactions of the Royal Society B: Biological Sciences 366:2351–2363.

4. Presley SJ, Higgins CL, Willig MR. 2010. A comprehensive framework for the evaluation of metacommunity structure. Oikos 119:908–917.

5. Chase JM, Kraft NJB, Smith KG, Vellend M, Inouye BD. 2011. Using null models to disentangle variation in community dissimilarity from variation in α-diversity. Ecosphere 2.

6. Ning D, Deng Y, Tiedje JM, Zhou J. 2019. A general framework for quantitatively assessing ecological stochasticity. Proceedings of the National Academy of Sciences of the United States of America 116:16892–16898.

7. Zhou J, Ning D. 2017. Stochastic Community Assembly: Does It Matter in Microbial Ecology? Microbiology and Molecular Biology Reviews 81.

8. Stegen JC, Lin X, Fredrickson JK, Chen X, Kennedy DW, Murray CJ, Rockhold ML, Konopka A. 2013. Quantifying community assembly processes and identifying features that impose them. The ISME Journal 7:2069–2079.

9. Tucker CM, Shoemaker LG, Davies KF, Nemergut DR, Melbourne BA. 2016. Differentiating between niche and neutral assembly in metacommunities using null models of ß-diversity. Oikos 125:778–789.

10. Lee SH, Sorensen JW, Grady KL, Tobin TC, Shade A. 2017. Divergent extremes but convergent recovery of bacterial and archaeal soil communities to an ongoing subterranean coal mine fire. ISME Journal 11:1447–1459.

11. Verster AJ, Borenstein E. 2018. Competitive lottery-based assembly of selected clades in the human gut microbiome. Microbiome 6:1–17.

12. Darcy JL, Washburne AD, Robeson MS, Prest T, Schmidt SK, Lozupone CA. 2020. A phylogenetic model for the recruitment of species into microbial communities and application to studies of the human microbiome. The ISME Journal 14:1359–1368.

13. Stegen JC, Lin X, Konopka AE, Fredrickson JK. 2012. Stochastic and deterministic assembly processes in subsurface microbial communities. ISME Journal 6:1653–1664.

14. Püttker T, de Arruda Bueno A, Prado PI, Pardini R. 2015. Ecological filtering or random extinction? Beta-diversity patterns and the importance of niche-based and neutral processes following habitat loss. Oikos 124:206–215.

15. Vellend M, Srivastava DS, Anderson KM, Brown CD, Jankowski JE, Kleynhans EJ, Kraft NJB, Letaw AD, Macdonald AAM, Maclean JE, Myers-Smith IH, Norris AR, Xue X. 2014. Assessing the relative importance of neutral stochasticity in ecological communities. Oikos 123:1420–1430.

16. Carroli IT, Cardinale Bradley J, Nisbet RM. 2011. Niche and fitness differences relate the maintenance of diversity to ecosystem function. Ecology 92:1157–1165.

17. Hassani MA, Özkurt E, Franzenburg S, Stukenbrock EH. 2020. Ecological Assembly Processes of the Bacterial and Fungal Microbiota of Wild and Domesticated Wheat Species. Phytobiomes Journal PBIOMES-01-20-0.

18. Sale PF. 1979. Recruitment, loss and coexistence in a guild of territorial coral reef fishes. Oecologia 42:159–177.

19. Verster AJ, Borenstein E. 2018. Competitive lottery-based assembly of selected clades in the human gut microbiome. Microbiome 6:186.

20. D’Andrea R, Riolo M, Ostling AM. 2019. Generalizing clusters of similar species as a signature of coexistence under competition. PLoS Computational Biology 15:1–19.

21. Nikolova C, Ijaz UZ, Magill C, Kleindienst S, Joye S, Gutierrez T. 2020. Response and oil degradation activities of a northeast Atlantic bacterial community to biogenic and synthetic surfactants. bioRxiv 2020.12.18.423525.

22. Leibold MA, Mikkelson GM. 2002. Coherence, species turnover, and boundary clumping: Elements of meta-community structure. Oikos 97:237–250.

23. Zhou J, Deng Y, Zhang P, Xue K, Liang Y, Van Nostrand JD, Yang Y, He Z, Wu L, Stahl DA, Hazen TC, Tiedje JM, Arkin AP. 2014. Stochasticity, succession, and environmental perturbations in a fluidic ecosystem. Proceedings of the National Academy of Sciences of the United States of America 111:E836–45.

24. Dini-Andreote F, Stegen JC, Van Elsas JD, Salles JF. 2015. Disentangling mechanisms that mediate the balance between stochastic and deterministic processes in microbial succession. Proceedings of the National Academy of Sciences of the United States of America 112:E1326–E1332.

25. Yakimov MM, Timmis KN, Golyshin PN. 2007. Obligate oil-degrading marine bacteria. Current Opinion in Biotechnology 18:257–266.

26. Hazen TC, Dubinsky EA, DeSantis TZ, Andersen GL, Piceno YM, Singh N, Jansson JK, Probst A, Borglin SE, Fortney JL, Stringfellow WT, Bill M, Conrad ME, Tom LM, Chavarria KL, Alusi TR, Lamendella R, Joyner DC, Spier C, Baelum J, Auer M, Zemla ML, Chakraborty R, Sonnenthal EL, D’haeseleer P, Holman H-YN, Osman S, Lu Z, Van Nostrand JD, Deng Y, Zhou J, Mason OU. 2010. Deep-Sea Oil Plume Enriches Indigenous Oil-Degrading Bacteria. Science 330:204–208.

27. Hamdan LJ, Fulmer PA. 2011. Effects of COREXIT EC9500A on bacteria from a beach oiled by the Deepwater Horizon spill. Aquatic Microbial Ecology 63: 101–109.

28. Gutierrez T, Biddle JF, Teske A, Aitken MD. 2015. Cultivation-dependent and cultivation-independent characterization of hydrocarbon-degrading bacteria in Guaymas Basin sediments. Frontiers in Microbiology 6:1–12.

29. Kleindienst S, Seidel M, Ziervogel K, Grim S, Loftis K, Harrison S, Malkin SY, Perkins MJ, Field J, Sogin ML, Dittmar T, Passow U, Medeiros PM, Joye SB. 2015. Chemical dispersants can suppress the activity of natural oil-degrading microorganisms. Proceedings of the National Academy of Sciences 112:14900–14905.

30. Liu J, Bacosa HP, Liu Z. 2017. Potential environmental factors affecting oil-degrading bacterial populations in deep and surface waters of the Northern Gulf of Mexico. Frontiers in Microbiology 7:1–14.

31. Miller JI, Techtmann S, Joyner D, Mahmoudi N, Fortney J, Fordyce JA, Garajayeva N, Askerov FS, Cravid C, Kuijper M, Pelz O, Hazen TC. 2020. Microbial communities across global marine basins show important compositional similarities by depth. mBio 11:1–12.

32. Kleindienst S, Grim S, Sogin M, Bracco A, Crespo-Medina M, Joye SB. 2016. Diverse, rare microbial taxa responded to the Deepwater Horizon deep-sea hydrocarbon plume. The ISME Journal 10:1–16.

33. Techtmann SM, Zhuang M, Campo P, Holder E, Elk M, Hazen TC, Conmy R, Domingo JWS. 2017. Corexit 9500 enhances oil biodegradation and changes active bacterial community structure of oilenriched microcosms. Applied and Environmental Microbiology 83:1–14.

34. Langenheder S, Ragnarsson H. 2007. The role of environmental and spatial factors for the composition of aquatic bacterial communities. Ecology 88:2154–2161.

35. Dumbrell AJ, Nelson M, Helgason T, Dytham C, Fitter AH. 2010. Relative roles of niche and neutral processes in structuring a soil microbial community. ISME Journal 4:337–345.

36. Stegen JC, Lin X, Fredrickson JK, Konopka AE. 2015. Estimating and mapping ecological processes influencing microbial community assembly. Frontiers in Microbiology 6:370.

37. Langenheder S, Wang J, Karjalainen SM, Laamanen TM, Tolonen KT, Vilmi A, Heino J. 2017. Bacterial metacommunity organization in a highly connected aquatic system. FEMS Microbiology Ecology 93:1–9.

38. Wu W, Lu HP, Sastri A, Yeh YC, Gong GC, Chou WC, Hsieh CH. 2018. Contrasting the relative importance of species sorting and dispersal limitation in shaping marine bacterial versus protist communities. ISME Journal 12:485–494.

39. Vass M, Székely AJ, Lindström ES, Langenheder S. 2020. Using null models to compare bacterial and microeukaryotic metacommunity assembly under shifting environmental conditions. Scientific Reports 10:1–13.

40. Evans S, Martiny JBH, Allison SD. 2017. Effects of dispersal and selection on stochastic assembly in microbial communities. ISME Journal 11:176–185.

41. R Core Team. 2019. R: A language and environment for statistical computing. 3.5.3. R Foundation for Statistical Computing, Vienna, Austria.

