## Supplementary Information for "Use of null models to compare the assembly of northeast Atlantic bacterial community in the presence of crude oil with either chemical dispersant or biosurfactant"

### Supplementary Methods

#### Elements of metacommunity structure (EMS)

EMS were assessed following the frameworks developed by Leibold and Mikkelson (1) and Presley et al. (2). For this, the ASV matrix was hierarchically analysed using three tests (coherence, turnover and boundary clumping). The R package *metacom* (3) was used to detect any pattern the metacommunity type responsible for the assembly structure of the microbial community. The outcome of coherence test, which counts the number of interruptions in species distributions in the ordinated matrix and comparing the empirical value to a null distribution, was the basis for the subsequent statistical analysis (2).

#### Incidence-based beta diversity

The incidence-based beta diversity (Raup-Crick) dissimilarity indices (ß_RC_) were calculated to test whether stochastic or deterministic processes were dominating the assembly of the microbial communities in the different treatments as described in Chase et al. (4). The function ‘raup-crick’ within the *vegan* package in R (5) was used to calculated the ß_RC_. The Raup-Crick metric was standardised to range from -1 to +1 and randomised 999 times. Values of ß_RC_ that were not statistically different from 0 indicate that the community is stochastically assembled. ß_RC_ values close to -1 or +1 indicate deterministically assembled communities that are more similar or dissimilar to each other than expected by chance, respectively.

#### Normalised stochasticity ratio (NST)

The NST for each treatment was calculated to measure the actual contribution of determinism in relation to stochasticity based on abundance-based similarity Ružička metric using null model algorithm PF as recommended by Ning et al. (6). For this, the ‘tNST’ function from *NST* package in R was used (6). When using abundance-based metric Ružička, null taxa abundances in each sample are calculated as random draw (1000 times) of the observed number of individuals with probability proportional to regional relative abundances of null taxa in the treatments. The microbial community assembly is completely deterministic when NST is 0% and completely stochastic when NST is 100%. Nonparametric permutational multivariate analysis of variance (PERMANOVA) was used to test whether the different treatments differed in their NST. Next, we calculated the NST of the oil-amended seawater (WAF), oil+dispersant amended (CEWAF), oil+biosurfactant (BEWAF), and non-amended seawater (SW) treatments over time to determine whether the proportion of stochasticity changed with time.

#### Tucker’s beta-null model

Beta-null modelling was performed to differentiate between neutral and niche assembly processes by implementing the framework developed by Tucker et al. (7) and Lee et al. (8), and, therefore, called Tucker’s beta-null model hereafter. Abundance-based (Bray-Curtis) and generalised UniFrac dissimilarity (abundance and phylogenetic information) metrics were used to calculate the beta-null deviation values, which are indicative of the magnitude of deviation between the observed and expected beta-diversity from randomly assembled pair of samples. The number of randomisations was set to 999. Beta-null deviation values were calculated only for the three oil amended treatments (WAF, CEWAF and BEWAF). The beta-null deviation values between treatments for each time point were tested for significance with two-way ANOVA and *post-hoc* Tukey’s test. Significance was accepted for p-value < 0.05.

#### Quantitative process estimates (QPE)

QPE require taxa to have different habitat associations (i.e. phylogenetic signal) so that phylogenetic turnover (the evolutionary distance differentiating taxa in one community from taxa in another community) can be estimated (9). To perform QPE, the extent to which the abundance-weighted ß-mean-nearest taxon distance (ßMNTD) deviated from the mean of the null distribution was determined (after 999 randomisations) and the significance was evaluated by the ß-Nearest Taxon Index (ßNTI; difference between observed ßMNTD and the mean of the null distribution in units of SDs). For instance, variable selection assembles the microbial community if the ßNTI value is greater than 2. In contrast, the community is assembled by homogeneous selection when the ßNTI is less than -2. In the case when there is no significant deviation from the null expectation, dispersal limitation, homogenising dispersal (mass effect) or random rift should drive the observed differences in phylogenetic community composition. To determine the relative importance of each of these processes, the abundance-based (Raup-Crick) beta-diversity was calculated using pairwise Bray-Curtis dissimilarity metric (ß_RCbray_) (9).

#### Lottery-based assembly model

The lottery-based assembly model was used to characterise the species distribution across treatments amended with crude oil with/without dispersant or rhamnolipid biosurfactant and to identify any microbial guilds whose distribution reflect a competitive lottery schema (10). The protocol developed by Verster and Borenstein (11) was followed with some exceptions. Briefly, the first step was to quantify how often species distribution within a group/guild includes a lottery winner (i.e., a group member that represent > 90% of the group’s abundance). The background expectation of the winner prevalence parameter was determined by implying a null model on the species abundances, which assumes a stick breaking process (12, 13). Second, a measure of the diversity of lottery winners was calculated by the Shannon diversity of the distribution of winners across samples. This measure is referred to as the frequency at which each ASV occurs as the lottery winner among all samples/treatments in which lottery winner is observed. ASVs that had less than 5,000 reads, appear at < 0.05% abundance, and had less than 3 ASVs were filtered out of the analysis.

#### Phylogenetic dispersion model

The aim of the phylogenetic dispersion model was to estimate the extent of recruitment of new species in a microbial community over time based on how similar or dissimilar they are from previously recruited species. For this, the null model of Darcy et al. (14) was applied. The model characterised the probabilities of detecting new species in a local community over time and then simulated the data 500 times to produce surrogate datasets forward in time, which were then used to evaluate the null polydispersity distribution *D*^ and the amount of polydispersity accumulated over time PD_m_. A logistical error model (14) is then performed to generate the dispersion parameter *D* which determined the extent to which either closely related or distant species were preferentially added to the surrogate community. *D* value > 0 means that phylogenetically distant species are preferentially recruited in the local community (overdispersion; phylogenetically divergent), whereas *D* < 0 indicate the opposite – phylogenetically similar species are detected in the local community (underdispersion; phylogenetically constrained). If *D* = 0, all species have the same probability of being detected for the first time (neutral).

**Supplementary Table S1**

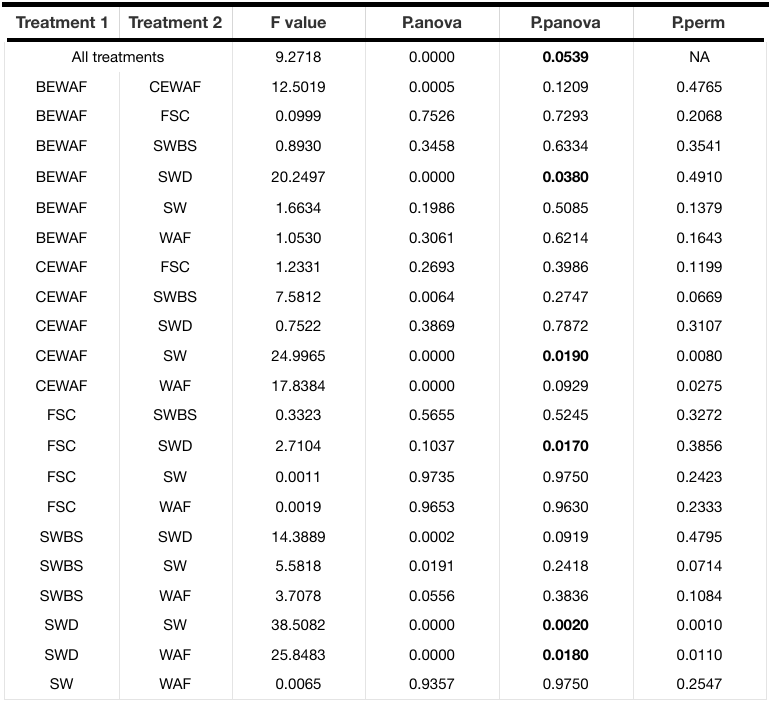

**Supplementary Figure S1.**

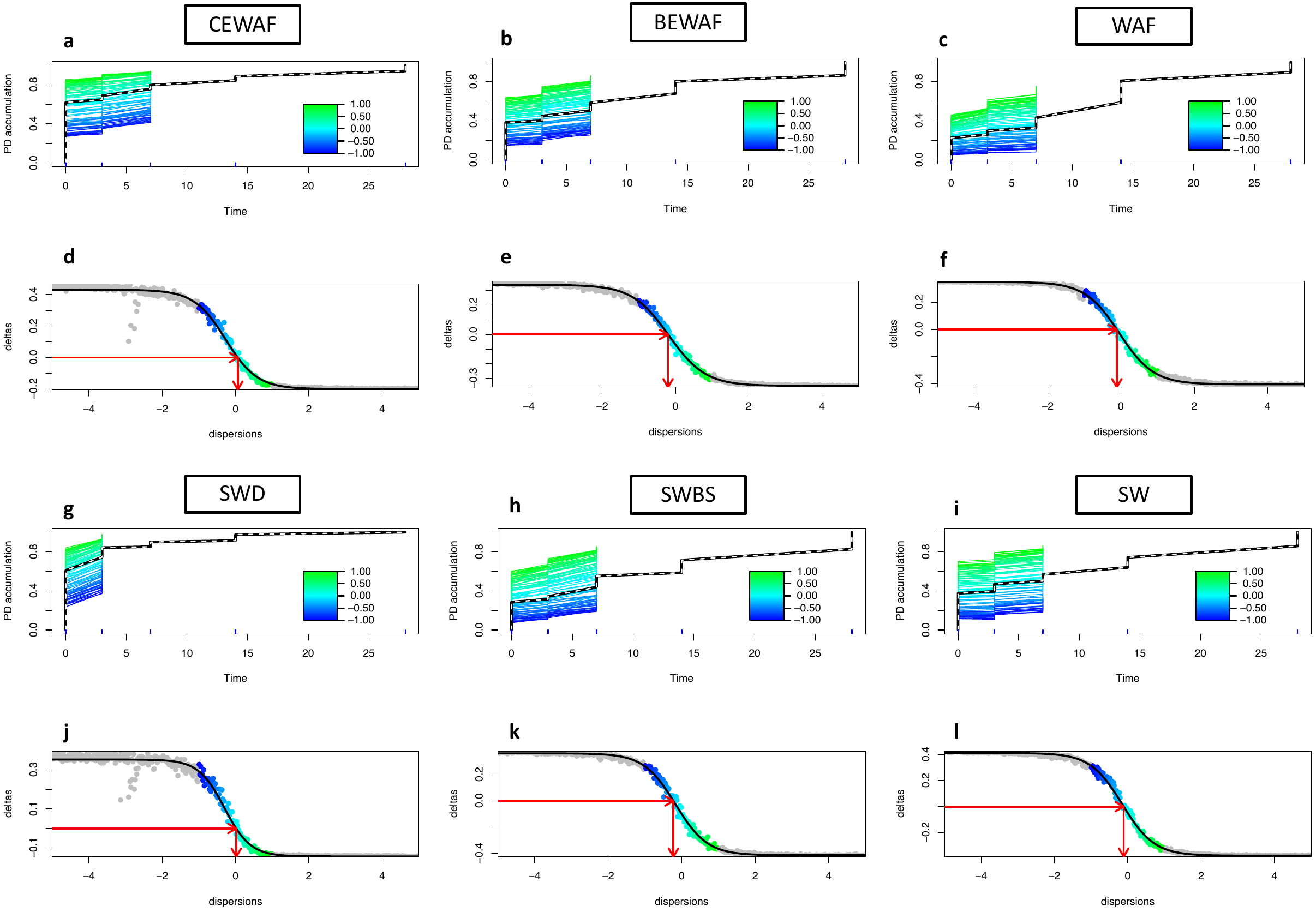

**References**

1.	Leibold MA, Mikkelson GM. 2002. Coherence, species turnover, and boundary clumping: Elements of meta-community structure. Oikos 97:237–250.

2.	Presley SJ, Higgins CL, Willig MR. 2010. A comprehensive framework for the evaluation of metacommunity structure. Oikos 119:908–917.

3.	Dallas T. 2014. Metacom: An R package for the analysis of metacommunity structure. Ecography 37:402–405.

4.	Chase JM, Kraft NJB, Smith KG, Vellend M, Inouye BD. 2011. Using null models to disentangle variation in community dissimilarity from variation in α-diversity. Ecosphere 2.

5.	Oksanen AJ, Blanchet FG, Friendly M, Kindt R, Legendre P, Mcglinn D, Minchin PR, Hara RBO, Simpson GL, Solymos P, Stevens MHH, Szoecs E. 2019. Community ecology package ‘ vegan ’. R package version 2.5-6.

6.	Ning D, Deng Y, Tiedje JM, Zhou J. 2019. A general framework for quantitatively assessing ecological stochasticity. Proceedings of the National Academy of Sciences of the United States of America 116:16892–16898.

7.	Tucker CM, Shoemaker LG, Davies KF, Nemergut DR, Melbourne BA. 2016. Differentiating between niche and neutral assembly in metacommunities using null models of β-diversity. Oikos 125:778–789.

8.	Lee SH, Sorensen JW, Grady KL, Tobin TC, Shade A. 2017. Divergent extremes but convergent recovery of bacterial and archaeal soil communities to an ongoing subterranean coal mine fire. ISME Journal 11:1447–1459.

9.	Stegen JC, Lin X, Fredrickson JK, Chen X, Kennedy DW, Murray CJ, Rockhold ML, Konopka A. 2013. Quantifying community assembly processes and identifying features that impose them. The ISME Journal 7:2069–2079.

10.	Sale PF. 1979. Recruitment, loss and coexistence in a guild of territorial coral reef fishes. Oecologia 42:159–177.

11.	Verster AJ, Borenstein E. 2018. Competitive lottery-based assembly of selected clades in the human gut microbiome. Microbiome 6:186.

12.	Macarthur RH. 1957. On the relative abundance of bird species. Proceedings of the National Academy of Sciences 43:293–295.

13.	Higgins CL, Strauss RE. 2008. Modeling Stream Fish Assemblages with Niche Apportionment Models: Patterns, Processes, and Scale Dependence. Transactions of the American Fisheries Society 137:696–706.

14.	Darcy JL, Washburne AD, Robeson MS, Prest T, Schmidt SK, Lozupone CA. 2020. A phylogenetic model for the recruitment of species into microbial communities and application to studies of the human microbiome. The ISME Journal 14:1359–1368.
